# A first global phylogeny of *Maize chlorotic mottle virus*

**DOI:** 10.1101/209940

**Authors:** Luke A Braidwood, Diego F Quito-Avila, Darlene Cabanas, Alberto Bressan, Anne Wangai, David C. Baulcombe

## Abstract

*Maize chlorotic mottle virus* has been rapidly spreading around the globe over the past decade. The interactions of *Maize chlorotic mottle virus* with potyviridae viruses causes an aggressive synergistic viral condition - maize lethal necrosis, which can cause total yield loss. Maize production in sub-Saharan Africa, where it is the most important cereal, is threatened by the arrival of maize lethal necrosis. We obtained *Maize chlorotic mottle virus* genome sequences from across East Africa and for the first time from Ecuador and Hawaii, and constructed a phylogeny which highlights the similarity of Chinese to African isolates, and Ecuadorian to Hawaiian isolates. We used a measure of clustering, the adjusted Rand index, to extract region-specific SNPs and coding variation that can be used for diagnostics. The population genetics analysis we performed shows that the majority of sequence diversity is partitioned between populations, with diversity extremely low within China and East Africa.

## Introduction

*Maize chlorotic mottle virus* (MCMV) is a positive-sense single-stranded RNA virus, and the sole member of the *Machlomovirus* genus in the tombusviridae family. The virus was described in Peru, then reported shortly afterwards in Brazil, Argentina and the USA in the 1970s^1–3^. Within the past decade, MCMV has spread globally, with first reports in China, Taiwan, Ecuador, Spain, and widely in East Africa (fig. 1a)^4–9^. MCMV can spread via soil water, seed, mechanical transmission, and is semi-persistently vectored by chrysomelid beetles and thrips (*Frankliniella*)^3, 7, 10–14^.

**Figure 1.**
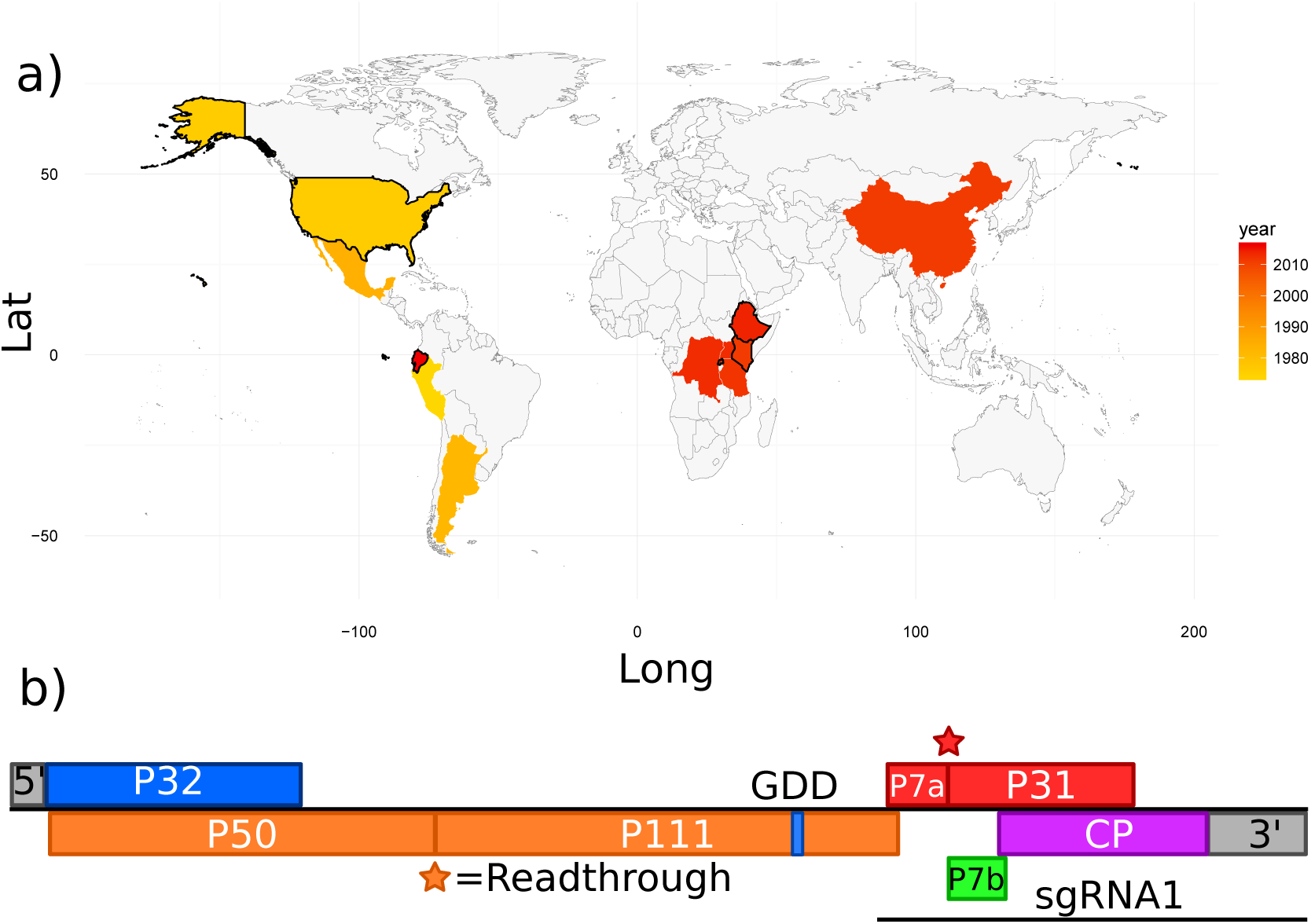
A) Global distribution of *Maize chlorotic mottle virus* (MCMV), coloured by year of first report. Countries sampled in this study are outlined in black. B) MCMV genome structure.

MCMV interacts synergistically with members of the potyviridae family: the potyviruses *Sugarcane mosaic virus* (SCMV) and *Maize dwarf mosaic virus* (MDMV), and the tritimovirus *Wheat streak mosaic virus* (WSMV). The interaction causes a more aggressive condition known as maize lethal necrosis (MLN). MLN causes around 80% yield loss in heavily affected areas, and destroyed an estimated 22% of Kenya’s 2013 maize crop^7, 15^. SCMV is the reported co-infecting virus in recent reports of MLN outbreaks in China, East Africa, and Ecuador^7, 8, 16^. The recent spread of MLN reflects the spread of MCMV since SCMV has been present in East Africa, China and South America for decades^17–19^.

In the absence of additional control measures or resistant varieties, it is likely that MLN will spread further. Ecological niche modelling suggests that large areas of Eastern sub-Saharan Africa are at high risk of M LN outbreaks^20^. Due to its rapid spread and interaction with local viruses, and the absence of resistant commercial maize lines, MCMV represents a significant threat to maize production in Sub-Saharan Africa, where it is the most important cereal crop. To investigate the spread of MCMV globally, and inform future control strategies, we decided to study genomic variation in MCMV isolates.

MCMV has a monopartite 4.4 kb genome (fig. 1b), which generates 1.4kb and 0.37kb sub-genomic RNAs during infection^21^. The genome encodes conserved tombusviridae proteins - the RNA-dependent RNA polymerase and associated protein (P111 and P50) and two movement proteins (P7a and P7b). MCMV expresses two unique proteins, P31 and P32, which have unknown function but are linked to systemic spread and viral accumulation respectively, and a viral suppressor of silencing (VSR) is yet to be identified^22^. P31 and P111 are expressed as readthrough proteins of P7a and P50 respectively, with P111 expressed through a conserved Tombusviridae amber stop codon readthrough motif (UAGGGR)^23^ (fig. 1b).

Full length MCMV genome sequences are available for North American, Chinese, and East African isolates, but until now there have been no data for South American and Hawaiian isolates. Here we use a combination of next-generation sequencing (NGS) and Sanger sequencing to obtain 37 novel MCMV genome sequences from East-Africa, Ecuador, and Hawaii, construct a first global phylogeny of MCMV, and identify genomic regions of interest for functional analysis and strain-typing. We also report evidence for recombination, and a natural nonsense mutation of P31 present in Hawaii and Ecuador.

## Results and Discussion

### Sequencing of MCMV isolates

Between 2012 and 2016, we collected MCMV-infected maize leaves and generated 37 novel MCMV genome sequences from Kenya (n=24), Ethiopia (n=5), Rwanda (n=4), USA (Hawaii, n=1), and Ecuador (n= 3) (supplemental table S1). African sequences were obtained via NGS, and the remainder through Sanger sequencing (see methods). Genbank accession numbers and isolate details are in Table S1. Sequence lengths ranged from 4399-4440bp; this is 99-100% coverage compared to the longest previously reported MCMV genome sequence. Including all Genbank isolates, nucleotide identity between genomes ranged from 100% to 96.55%. Nucleotide diversity between all sequences was 0.01, which is very low for viruses generally and similar to the lowest recorded diversity values in the Tombusviridae (table 1)^24–26^.

**Table 1.**
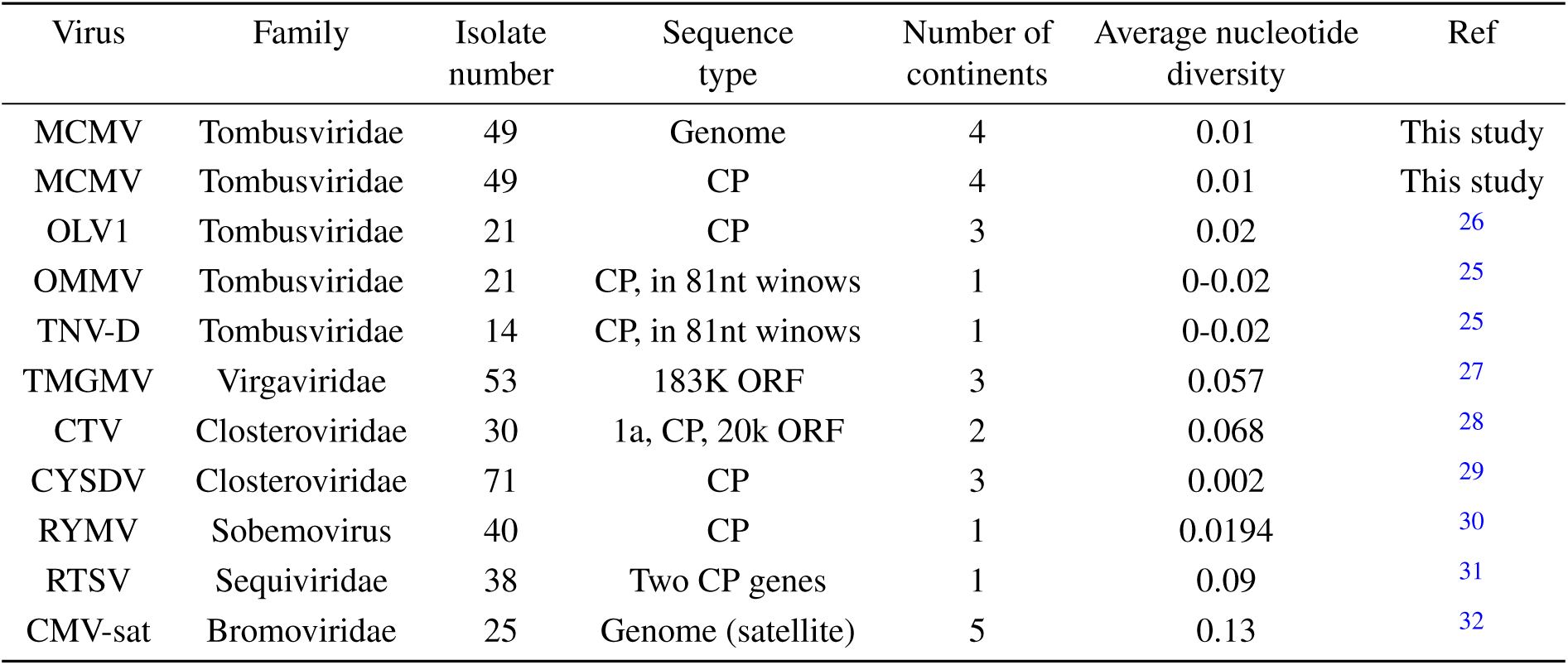
Comparison of average nucleotide diversities calculated for positive-sense ssRNA viruses. CMV-sat=*Cucumber mosaic virus satellite RNA*, CTV=*Citrus tristeza virus*, CYSDV=*Cucurbit yellow stunting disorder virus*, MCMV=*Maize chlorotic mottle virus*, OLV1=*Olive latent virus-1*, OMMV=*Olive mild mosaic virus*, RTSV=*Rice tungro spherical virus*, RYMV=*Rice yellow mottle virus*, TNV-D=*Tobacco necrosis virus D*, TMGMV=*Tobacco mild green mosaic virus*.

### Phylogenetic analysis

Phylogenetic analyses assume a single evolutionary history of each genome, an assumption which may be violated in the case of extensive recombination. We therefore performed a splits network analysis, which detects conflicting phylogenetic signals that can be caused by recombination. There is clear separation of isolates from different regions, indicating that there has been no recombination between MCMV genomes in geographically isolated regions (fig. 2a). Conventional phylogenetic analysis is suitable, therefore, for investigating the relationships between regions. To construct an unrooted phylogenetic tree, we used a nucleotide alignment containing all novel MCMV isolates, including ambiguous bases at the 10% threshold, and genome sequences available in GenBank. We chose nucleotide alignment to enhance resolution due to the low divergence between MCMV isolates, and used Bayesian inference (MrBayes 3.2) to generate the tree (fig. 2b). Clearly resolved clades contain North American isolates, Hawaiian with South American isolates, and Chinese with African isolates.

**Figure 2.**
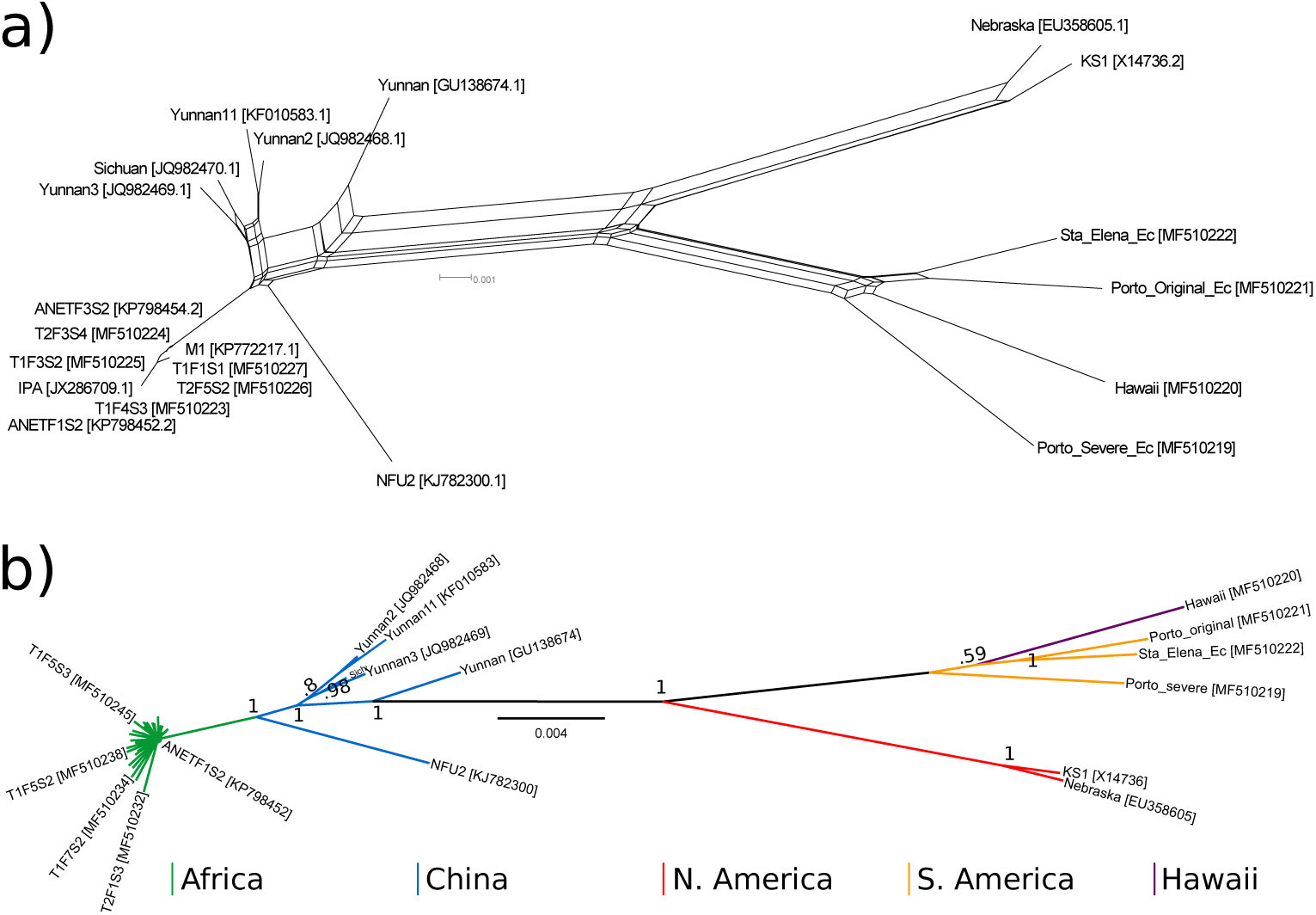
A) Splits network of *Maize chlorotic mottle virus* (MCMV) genomes, distances calculated with uncorrected P, and network generated by neighbour-net in Splitstree V4.6. B) MCMV phylogeny generated using Bayesian inference in Mr Bayes 3.2. Scale bars are nucleotide substitutions per site.

The phylogenetic tree shows clear separation between geographic regions, with the exception of Hawaii and Ecuador. Phylogenetic confidence estimates (fig. 2b) and population genetics (below) illustrate that the majority of MCMV genome diversity occurs between sub-populations. For China, East Africa, and Ecuador, intra-group diversity is minimal, suggesting either (separate) single introduction events or repeated introductions from the same sub-populations. The similarity of Ecuadorian sequences to the Hawaiian sequence, and African to Chinese sequences, could represent the epidemiological routes of MCMV across the globe; typically the origin of an epidemic is identified as the most closely related non-epidemic strain^33^. In this case, that would make China the source of the East African outbreak, and Hawaii the source of the Ecuadorian; however corroborating evidence and more complete sampling of global MCMV genomes would be required to confirm this. In particular, sequence data from other Central and South American regions with MCMV presence could determine whether the MCMV outbreak in Ecuador is linked most closely to neighbouring countries or Hawaii. It would also determine whether the observed limited sequence divergence amongst MCMV isolates is universal, suggesting either recent evolution or intense purifying selection, or whether currently undocumented diversity is present in its presumed ancestral range – the ancestral ranges of *Teosinte* and maize.

The phylogenetic proximity of Ecuadorian and Hawaiian isolates could be explained by seed transmission. Seed transmission of MCMV has been reported at a rate of 0-0.33%^12^. This rate may be an underestimate as parent populations were naturally infected (so likely had *<*100% infection). 12 of 26 ten-seed samples bought at Kenyan markets and tested by RT-PCR were positive for MCMV, and 72% of kernels from a single plant, which demonstrates MCMV presence in commercial maize seed, but not transmission^7^. Seeds from two commercial hybrids planted in the region of the 2015 Ecuadorian outbreak were grown in sterile soil inside insect-proof growth chambers, and 8 and 12% of seedlings were MCMV-infected^8^. These isolates, and the Ecuadorian genomes collected in this study, were most closely related to the Hawaiian genome. Hawaii is an important region for maize seed production for a number of major agricultural companies, although it is unknown whether the infected seeds from Ecuador were produced in Hawaii. Regardless, it is vital that the ability of MCMV to be transmitted via maize seed is investigated further, especially given Hawaii’s central role in maize seed production and the rapid spread of MCMV across East Africa. However, given the low rate of seed transmission, and the high populations of thrip vectors in the *Frankliniella* genus, we would suggest although seed transmission may be important for long distance dispersal, once arriving in an area vector and soil transmission will likely be more important for local spread^34, 35^.

### SNP and coding analysis

Structural variation within whole genomes was limited to the genomic termini. We found small 5’ extensions of the genome – African isolates and the Taiwanese sweetcorn isolate (KJ782300.1) have a G insertion, giving a 5’ terminal AGGG, as opposed to the AGG found in isolates from other regions. The G9 sequence at the 3’ terminus of the genome has a G insertion in North American isolate X14736.2, while the first reported Kenyan isolate, JX286709.1, has a G insertion immediately downstream of the TAAT sequence in the 3’ terminus (fig. 3a and b).

**Figure 3.**
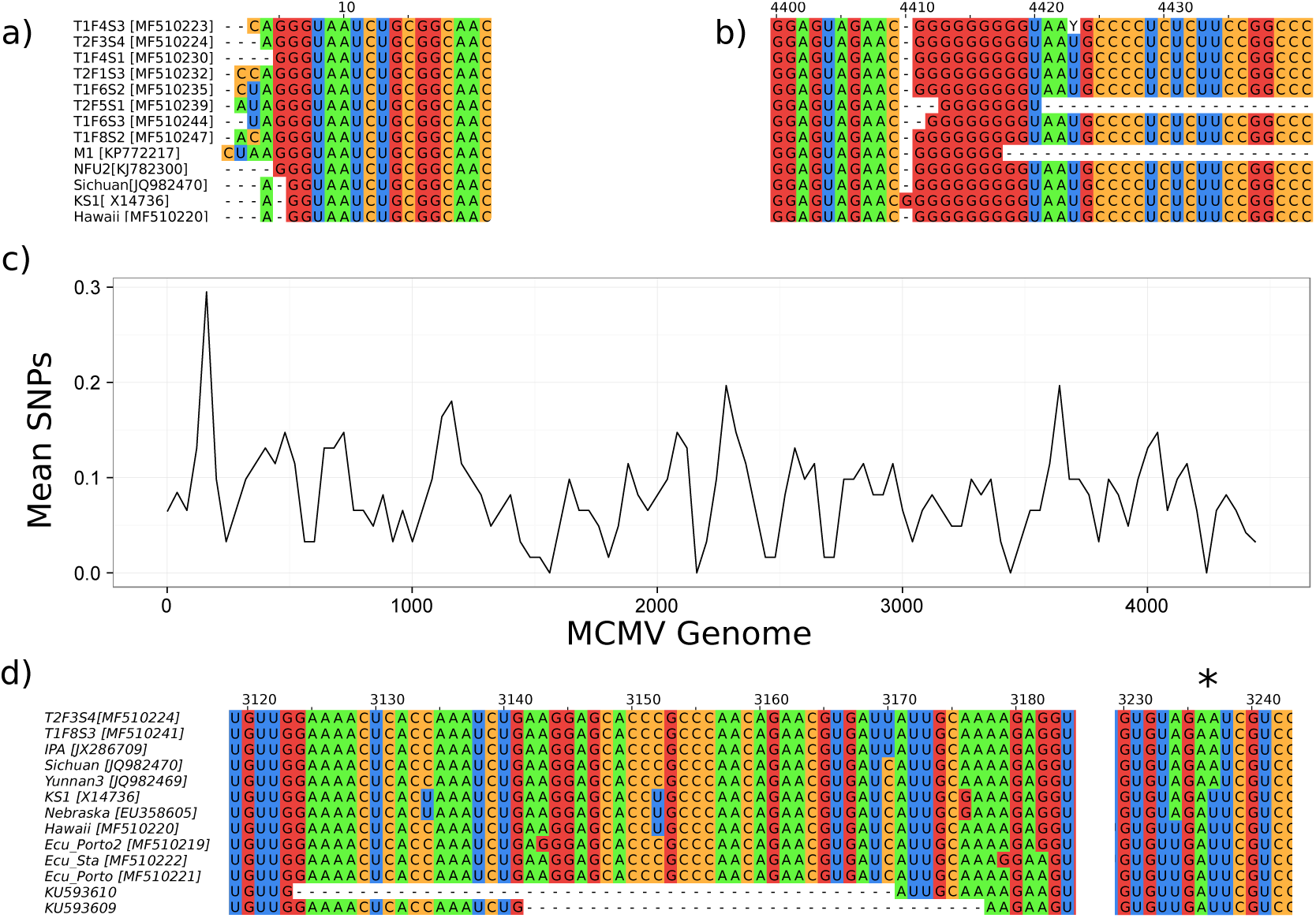
Genomic termini of the *Maize chlorotic mottle virus* (MCMV) genome, 5’ (A) and 3’ (B). Sequences with identical termini have been removed for clarity. C) SNP distribution across the MCMV genome, in 60bp windows calculated every 40bp. D) Variation in P31, with early stop codon indicated by asterisk.

There were 419 mutations at 388 polymorphic sites across the MCMV genome in our dataset (fig. 3c). To identify geographic region-specific genomic variation we used the Adjusted Rand Index (ARI). The ARI is a chance-adjusted measure of the similarity between two data clusterings, varying between −1 (dissimilar) and 1 (similar). We used the phylogenetic clades (Africa, China, North America, and South America with Hawaii) as one data cluster, and a site in the nucleotide alignment for the other. Therefore, ARI values above zero were produced when SNPs segregated according to the phylogenetic clades. To identify sites which clustered significantly better than chance, we randomised the members of each geographic clade and recorded the ARI across the genome 1000 times. Sites above the 95% level of the randomised ARI scores were taken to be significant. Significant sites were extracted, identifying candidate SNPs for (uncharacterised) phenotypic variation between clades, which can be used to design diagnostics appropriate for separating isolates from different regions (supplemental table S3).

To identify protein coding changes within the clades we repeated our ARI analysis on translated alignments for each MCMV ORF (supplemental table S4-S9). The most significant variation was observed in the coat protein gene and the P7a/P31 region in which the systemic movement protein P31 is expressed by readthrough of the P7a ORF UGA stop codon. We observed unreported variation in an exposed loop of the MCMV capsid - Phe76Leu in Chinese sugarcane (KF010583.1). Exposed capsid loops can function in vector transmission, and Wang *et al.* (2015) previously identified two variable residues in the exposed loops of the MCMV capsid, Pro81Ser in Nebraska (NC 003627) and Kansas (EU358605.1) isolates, and Ala62Asp in the 2012 Kenyan isolate (JX286709.1)^5, 36, 37^.

P31 enhances systemic MCMV movement via an unknown mechanism^22^. However, in Ecuadorian and Hawaiian isolates there is an early stop codon 18bp downstream of the P7a UGA stop codon, truncating the majority of P31 (fig. 3d). This unexpected stop codon is also present in the two previously reported partially sequenced Ecuadorian MCMV isolates, in which there are also 47 and 36bp deletions in the P7a and P31 coding sequences upstream of the stop codon^8^. MCMV P31 has unknown function, a unique carboxy-terminal extension, and mutagenesis experiments suggest that it promotes systemic movement (Scheets, 2016). Both the Hawaiian isolate and Ecuadorian isolates contain an early stop codon, truncating P31 six amino acids after the read-through stop codon of the movement protein P7a. This is unlikely to be a sequencing error, as *>*5 independent clones were sequenced in Hawaii and Ecuador for each isolate. The stop codon leaves 162nt of non-coding RNA in the Hawaiian and Ecuadorian isolates, before the capsid gene initiates. Interestingly, the partial Ecuadorian sequences reported previously also contain this stop codon, as well as deletions upstream within the P7a coding region^8^, which could represent selection pressure for smaller genomes once the function of P31 was lost, which is commonly observed in RNA viruses^38^.

Scheets (2016) used a mutant cDNA clone with an early stop codon mutation (p7bQ12N) to investigate P31 function, similar to the natural mutants we report from Hawaii and Ecuador^22^. Inoculation with p7bQ12N resulted in slower systemic spread of MCMV, and sequencing showed that the systemically infected leaves contained either a mixture of p7bQ12N with pseudorevertant (the stop codon removed), or exclusively pseudorevertant genomes. Mixtures were predominantly p7bQ12N, raising the possibility that the wild isolates from this study contained low-frequency genomes which complement the predominant genomes with mutated P31. Complementation of this form has been observed in tomato aspermy cucumovirus - 76% of genomes had a mutated movement protein, which were complemented by a minority of wild-type viral genomes within the host^39^. Experimentally mixing mutant and wild-type MCMV transcripts could establish whether this is a possibility for MCMV.

### Population genetics analysis

We examined diversity and population structure with all novel and publicly available MCMV sequences. Overall sequence diversity was low, with 45.5 nucleotide differences between sequences on average (table 2). Haplotype diversity was high, 0.99, with 42/49 haplotypes unique, although these are separated by a low number of SNPs. For subpopulation analysis, China and Africa were considered separately to allow estimation of gene flow between the populations. In terms of nucleotide diversity, the most diverse clade was South American and Hawaiian (pi = 0.015), while the least was Africa (pi = 0.0017), with only 7.51 nucleotide differences between African sequences on average. Variation in nucleotide identity was extremely limited for both Chinese and African sequences, with *>*99% sequence identity within each group. This high level of similarity suggests single introduction events for both the Chinese and East African outbreaks of MCMV.

**Table 2.** *Maize chlorotic mottle virus* genome diversity globally and within phylogenetic clades.

| Population | Genomes | Segregating sites | No. of haplotypes | Haplotype diversity | Average no. of SNPs | Nucleotide diversity |
| --- | --- | --- | --- | --- | --- | --- |
| Global | 49 | 388 | 42 | 0.986 | 45.5 | 0.0104 |
| South America and Hawaii | 4 | 123 | 4 | 1 | 64.7 | 0.0148 |
| North America | 2 | 22 | 2 | 1 | 22 | 0.00505 |
| China | 6 | 98 | 6 | 1 | 36.1 | 0.00829 |
| Africa | 37 | 114 | 30 | 0.974 | 7.51 | 0.00172 |

To test genetic differentiation between subpopulations we used Hudson’s test statistics (K_st_, K_st_*, Z, Z*, S_nn_), with all statistics indicating significant differentiation (table 3)^40, 41^. S_nn_ is powerful at all sample sizes and diversities, and so is most appropriate in this case, due to uneven sample sizes^41^. Hudson’s H_st_ tests differentiation based on haplotype diversity, and was not significant, presumably due to the high proportion of unique haplotypes across all populations. Likewise, F_st_ and N_st_ values of 0.74 indicate that a high proportion of the genetic variation in MCMV is explained by population structure^42, 43^. F_st_ and N_st_ values generated from pairwise comparisons between populations (table 4) illustrate that the sub-populations with least genetic variation explained by population structure are China and Africa, which is expected given their phylogenetic proximity.

**Table 3.** *Maize chlorotic mottle virus* population differentiation

| Population statistic | Data type | Value | P-value |
| --- | --- | --- | --- |
| H<sub>s</sub> | Haplotype | 0.978 | 0.072 |
| K<sub>s</sub> | Nucleotide | 16.3 | 0 |
| K<sub>s</sub>* | Nucleotide | 2.24 | 0 |
| Z | Nucleotide | 400 | 0 |
| Z* | Nucleotide | 5.69 | 0 |
| S<sub>nn</sub> | Nucleotide | 1 | 0 |

**Table 4.** *Maize chlorotic mottle virus* subpopulation differentiation

| Population 1 | Population 2 | H<sub>s</sub> | K<sub>s</sub> | N<sub>st</sub> | F<sub>st</sub> |
| --- | --- | --- | --- | --- | --- |
| South America and Hawaii | North America | 1 | 50.4 | 0.684 | 0.680 |
| South America and Hawaii | China | 1 | 47.5 | 0.638 | 0.633 |
| South America and Hawaii | Africa | 0.976 | 13.1 | 0.747 | 0.744 |
| North America | China | 1 | 32.6 | 0.790 | 0.786 |
| North America | Africa | 0.974 | 8.25 | 0.900 | 0.898 |
| China | Africa | 0.977 | 11.5 | 0.511 | 0.511 |

The HYPHY server (www.datamonkey.org) was used to test for selection using alignments corresponding to each ORF in the MCMV genome. MEME was used to detect episodic positive selection, with significant (p*<*0.01) codons found in P50, P32, P31, and P111 (supplemental table S2)^44^. IFEL was used for detection of positive and negative selection in each MCMV ORF (Tables S2). Significant (p*<*0.01) positive selection on codons was detected in P50, P32, and P31, with significant negative (purifying) selection found at sites in all ORFs (supplemental table S2). Examining the density of negatively selected sites across the genome showed that the regions under most intense purifying selection corresponded to the readthrough region of P111, and the C-terminus of the coat protein.

Unlike previous dN/dS-based analyses of MCMV, we detected significant positive selection in the genome: at sites in P50, P32, and P31, three proteins of unknown function^45^. This can be attributed to the greatly increased volume of sequence data included in the analysis. The positive selection may represent adaptation to specific host interactions across different maize lines, or to different environments. We detected purifying selection across the MCMV genome, especially in the post-readthrough region of the RdRP, which is highly conserved amongst Tombusviridae members, and the C-terminus of the CP. This is the location of an asymmetric unit important for interaction between capsid monomers in the assembled viral coat^46^. dN/dS analyses of selection in viral genomes can be complicated by selection at the RNA sequence level to preserve functional RNA structures, such as the Tombusviridae subgenomic RNA promoters or 3’Cap Independent Translational Enhancers^47^.

## Methods

### NGS of East African MCMV isolates

During August 2014 maize leaf samples were collected from Kenya and Ethiopia, stored on dry ice in RNA-later (Ambion), then RNA was extracted using Trizol (Ambion) according to manufacturer’s instructions. An additional Rwandan maize leaf RNA sample was received from FERA (UK). Ribosomal RNA (rRNA) depletion was performed with Ribo-Zero Magnetic Kit (Plant Leaf – Epicentre). Indexed stranded libraries were constructed using Scriptseq V2 RNA-Seq Library Preparation kits and Scriptseq Index PCR primers (Epicentre). Purification steps were performed using Agencourt AMPure XP beads. Library quantity and quality were checked using Qubit (Life Technologies) and on a Bioanalyzer High Sensitivity DNA Chip (Agilent Technologies). Libraries were sent to Beijing Genomics Institute for 100bp paired-end sequencing on one lane of a HiSeq 2000 (Illumina).

### NGS sequence assembly

Libraries were demultiplexed allowing one error within the index sequence using a custom python script, then adaptors trimmed using Trim galore! (parameters: –phred64 –fastQC –illumina –length 30 –paired –retain unpaired input 1.fq input 2.fq)^48, 49^.

Deduplication was performed by string-matching using scripts from the Quality Assessment of Short Read (QUASR) pipeline^50^. A bowtie reference was constructed from a fasta containing all 35 publicly available MCMV sequences using the bowtie2-build command, producing an index with each MCMV sequence as a separate chromosome. Reads were aligned to this index in order to capture all MCMV reads using bowtie2 (parameters: -D 20 -R 2 -N 1 -L 20 -i S,0,2.50 –phred64 –maxins 1000 –fr)^51^. Reads aligning to MCMV were assembled using Trinity, MCMV contigs of 1kb or more were extracted by blast, and contigs were manually checked and assembled if necessary^52^. Libraries were then realigned to their respective trinity contigs by bowtie2, pileups generated by samtools, and consensus sequences called using QUASR script pileup consensus.py, with a threshold of zero or ten % of reads to use ambiguity codes (parameters: –ambiguity 2—10 –dependent –cutoff 25 –lowcoverage 20)^53^.

### Sanger sequencing of Hawaiian and Ecuadorian samples

Hawaiian MCMV was purified from infected maize leaves, RNA extracted using Trizol, and RT-PCR performed. Primers based on those used to sequence the Nebraska isolate (EU358605.1) were used to amplify six amplicons covering the genome^54^. Amplicons were cloned into pCR4-TOPO TA plasmids and Sanger sequenced using M13 and internal primers to obtain total coverage. A consensus sequence was determined by sequencing six full-length clones.

A total of three Ecuadorean isolates were fully sequenced for this study. Primers used to amplify the genomes were designed from conserved regions using an alignment of available sequences including a local isolate obtained previously by degenerate-oligonucleotide-primed RT-PCR using double-stranded RNA as template^8^. Overlapping PCR amplification products (three independent reactions for each primer set) were sequenced in both directions by the Sanger method (Macrogen, South Korea).

### Sequence alignment

MCMV genome sequences were aligned using MUSCLE in MEGA6, with a gap extension cost of −1000. The alignment was checked and refined manually in JALview^55, 56^.

### Recombination Analysis

SplitsTree4 was used to generate splits networks, using the default settings - distances were calculated by uncorrected P (match option for ambiguous bases), and network generated by neighbour-net^57^. RDP4 was used to examine the evidence for recombination, utilising the algorithms RDP, GENECONV, Chimeara, MaxChi, BOOTSCAN and SISCAN^58^. A number of highly similar (*>*99.5% identity) African sequences were excluded from the analysis to make figures clearer and decrease computational time in RDP4.

### Phylogenetic Tree Construction

Alignment sites were split into three partitions: A) non-coding B) codon positions one and two and overlapping ORFs C) codon position three. To generate phylogenetic trees two runs of four Monte Carlo Markov Chain (MCMC) computations were run for 1000,000 generations under a general-time-reversible (GTR) model with a gamma distribution of rate variation between sites in MrBayes 3.2 ^59^. The first 10% of generations were discarded as burn-in (default). Convergence and effective sample size were examined using Tracer to confirm that estimated sample sizes for each parameter exceeded 200, as recommended by the MrBayes manual. We pooled together 1800 trees sampled every 500 generations (default) and constructed consensus trees.

### Population genetics and SNP analysis

To extract alignment sites at which data clustered similarly to phylogeny, we used the Adjusted Rand Index to measure the clustering between phylogenetic groupings and alignment sites, extracting those with a score above zero. This was performed using custom R scripts. Population genetics indices and statistics were produced using an alignment of MCMV sequences without ambiguity codes (i.e. 0% threshold) in DnaSP v5, as DnaSP v5 does not accept ambiguity codes^60^. Significance for test statistics was assessed by a permutation test with 1000 replications. To extract alignment sites with data clustering similarly to phylogeny, we used the Adjusted Rand Index to compare the phylogenetic groupings with alignment sites, extracting those with a score above zero.

## Acknowledgements

The authors would like to acknowledge colleagues at KALRO for their assistance in maize sampling, to farmers in all study areas for providing maize samples, to John Welch for guidance on phylogenetic analyses, and to Shizu Watanabe for her work on purifying MCMV in Hawaii. L.B. is supported by the BBSRC DTP, and D.C.B. is supported by the Royal Society Edward Penley Abraham Research Professorship.

## Author contributions statement

L.B. designed and performed experiments, analysed the results and wrote the manuscript, D.Q.A., D.C., and A.B. performed experiments, A.W. and D.C.B designed experiments. All authors reviewed the manuscript.

## Additional information

Novel MCMV genome sequences were deposited in Genbank, with the accession numbers MF510219-MF510251.

